# Development and validation of a miniaturized host range screening assay for bacteriophages

**DOI:** 10.1101/2021.02.18.431904

**Authors:** Renee Nicole Ng, Lucinda Jean Grey, Andrew Vaitekenas, Samantha Abagail McLean, Daniel Rodolfo Laucirica, Matthew Wee-Peng Poh, Jessica Hillas, Scott Glenn Winslow, Joshua James Iszatt, Thomas Iosifidis, Anna Sze Tai, Patricia Agudelo-Romero, Barbara Jane Chang, Stephen Michael Stick, Anthony Kicic

**Affiliations:** School of Biomedical Sciences, The University of Western Australia; (R.N.N.); (L.J.G.); Wal-Yan Respiratory Research Center, Telethon Kids Institute, Australia; (P.A.R.); (S.M.S.); (JH); (S.A.M); (S.W); (J.J.I.) (T.I.); Medical School, The University of Western Australia; (D.R.L.); (W.W-P.P); Occupation and the Environment, School of Public Health and Psychology, Curtin University, Australia; (AV); (A.K.); Department of Respiratory Medicine, Sir Charles Gairdner Hospital, Perth, WA, Australia; (A.S.T.); Institute for Respiratory Health, School of Medicine, The University of Western Australia, Perth, WA, Australia; The Marshall Center for Infectious Diseases Research and Training, School of Biomedical Sciences, The University of Western Australia; (B.J.C.); Department of Respiratory and Sleep Medicine, Perth Children’s Hospital, Australia; Centre for Cell Therapy and Regenerative Medicine, School of Medicine and Pharmacology, The University of Western Australia and Harry Perkins Institute of Medical Research, Australia

**Keywords:** bacteriophages, phage therapy, host range

## Abstract

Antimicrobial resistance is a global health crisis, partly contributed by inappropriate use of antibiotics. The increasing emergence of multidrug resistant infections has led to the resurgent interest in bacteriophages as an alternative treatment. Current procedures assessing susceptibility and breadth of host range to bacteriophage are conducted using large-scale manual processes that are labor-intensive. The aim here was to establish and validate a scaled down methodology for high-throughput screening in order to reduce procedural footprint. Bacteriophages were isolated from wastewater samples and screened for specificity against 29 clinical *Pseudomonas aeruginosa* isolates and PA01 using a spot test (2 μL/ drop). Host range assessment was performed on four representative *P. aeruginosa* isolates using both double agar overlay assay on petri dishes and 24-well culture plates. The breadth of host range of bacteriophages that exhibited lytic activity on *P. aeruginosa* isolates were corroborated between the current standard practice of whole plate phage assay and 24-well phage assay. The high correlation achieved in this study confirms miniaturization as the first step in future automation that could test phage diversity and efficacy as antimicrobials.

## 1. Introduction

Antimicrobial resistance (AMR) to both commonly used and last line antibiotics is increasing and adds an extra burden to healthcare systems [1–4]. Treatment is particularly challenging in those with chronic lung diseases such as cystic fibrosis (CF) and bronchiectasis where airways become chronically infected by pathogens such as *Pseudomonas aeruginosa* (*P. aeruginosa*) [5–7]. With little investment into the development of new antibiotics to counteract the issue of multidrug resistant (MDR) infections, alternative treatments are being sought. Bacteriophage (phage) therapy has been identified as a potentially useful strategy [8,9,18–20,10–17]. While it has a history of use in Eastern Europe [21–26] for decades, interest in the West was eclipsed with the advent of effective antibiotics [27–29].

A major obstacle in translating phage therapy into standard clinical practice, is that most in vitro and in vivo validations still need to be performed by academic research laboratories [8,10,12]. Further complications arise due to a lack of standardized procedures in research settings for the screening and selection of the phages for therapeutic use, which contrasts with current diagnostic practices in clinical and industrial laboratories. Academic research laboratories typically depend on manual manipulation of resources and labor, creating an “automation gap” [30]. This phenomenon does not exist in a clinical laboratory due to the necessity of a fast turnaround time (TAT), with the delivery of accurate results guiding treatments and improving clinical outcomes. One area in laboratory medicine: clinical chemistry, was among the first to implement automation, and observed a significant increase in both productivity and reduction of operational costs [31]. Turnaround time remains a benchmark and performance indicator of any current clinical/diagnostic laboratory audited by appropriate regulatory bodies [32].

Microbiological diagnoses have one of the slowest reported TAT. Many laboratories still engage in manual processes since they receive a diversity of sample types, however, the incubation time needed for traditional bacteria culture growth remains a significant contributor [33,34]. In the context of phage therapy and other associated therapeutic pipelines, this has a significant impact since the isolation and purification of phage are laborious and has a high procedural footprint in the laboratory. Furthermore, testing of host range via the traditional double agar overlay method is both relatively time-consuming and inefficient. Furthermore, the screening of phage efficacy also requires a vast number of laboratory consumables that can escalate exponentially with increasing number of phages and bacterial strains tested. Here we described a miniaturized host range screening assay against clinical isolates of P. aeruginosa developed in our laboratory to improve productivity and reduce procedural footprint.

## 2. Results

### 2.1 Isolation and purification of bacteriophages

A panel of 30 *P. aeruginosa* isolates (29 clinical isolates and PA01) was used as propagating hosts for the isolation and purification of phages. Wastewater samples were filtered through 0.22 μm bottle-top filters to remove debris and microorganisms prior to the isolation of phages. Positive phage activity was observed as a zone of clearing on a streak of a single *P. aeruginosa* isolate after 24 hours enrichment of filtered wastewater with a single isolate of *P. aeruginosa* as a zone of clearing (Figure 1a, left panel). Whole plate agar overlay of the enriched wastewater with *P. aeruginosa* showed plaques of different morphologies. A single plaque was picked and filtered for purification of phages (Figure 1b, right panel). Phages were purified over three rounds using whole plate agar overlay. Plaques retained similar morphologies through purification rounds (Figure 1b). Observed plaques included clear (Figure 1c; 1), clear with an uneven hazy ring (Figure 1c; 2), clear with bullseye (Figure 1c; 3), opaque with bullseye (Figure 1c; 4) and opaque (Figure 1c; 5) morphologies, indicating diversity in phage populations from waste water samples. A total of 231 phages were isolated after three rounds of purification.

**Figure 1.**
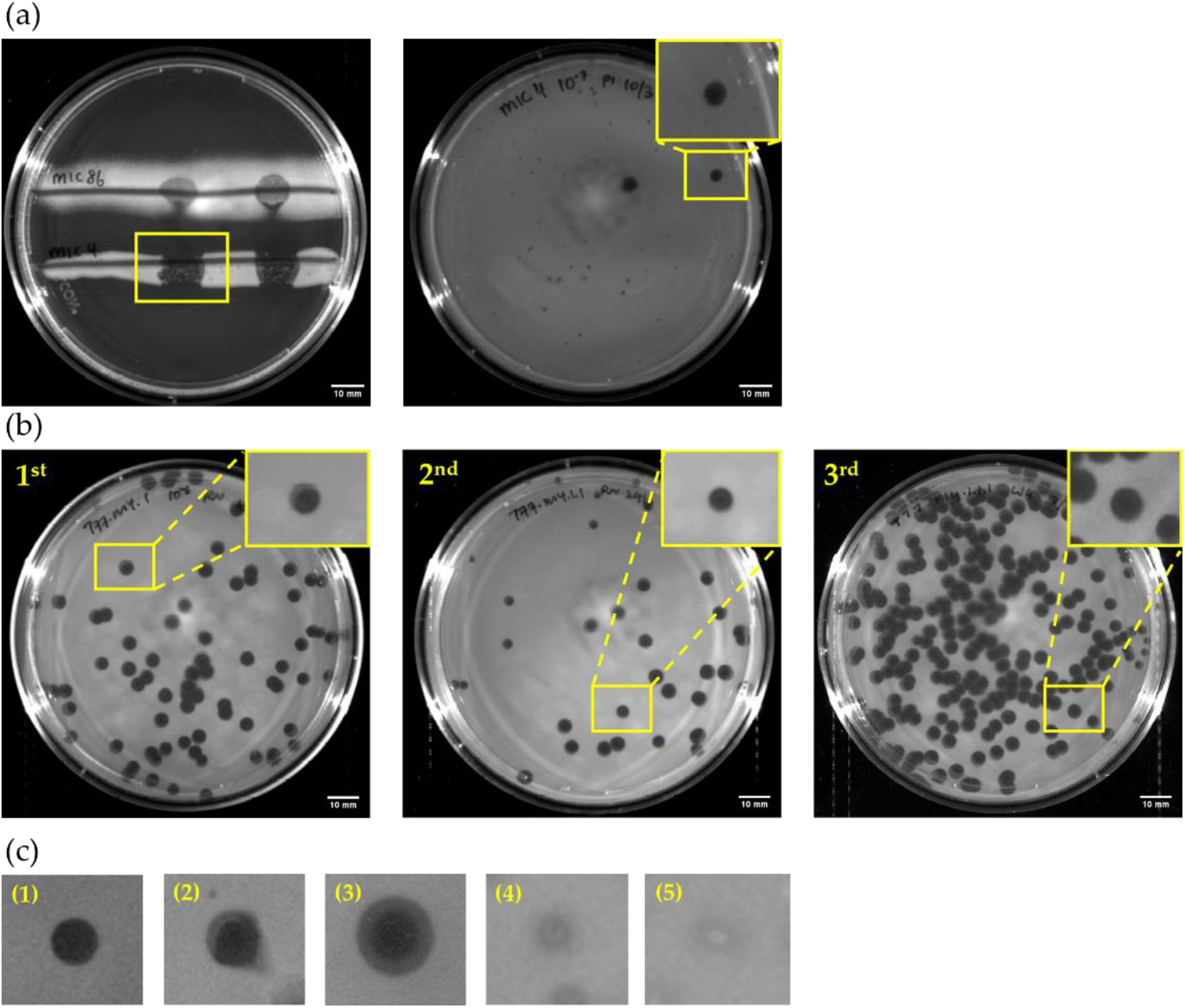
Phage isolation and purification. (a) Phage activity was tested against P. aeruginosa isolate (left panel). Whole plate agar overlay was performed to observe different morphologies present in the enriched wastewater sample. (b) Phages were selected and purified through three rounds of purification, plaque morphology was observed to exhibit similar zones of clearing. (c) Various plaque morphologies were observed from the whole plate agar overlay assay with the enriched wastewater samples.

### 2.2. Optimization of host range screening using miniaturized phage assay

To optimize the amount of Luria-Bertani Lennox (LB) overlay agar required per well, a range of agar volumes were tested and visually inspected the next day following an overnight static incubation at 37°C (Supplementary Table 1). Overlay agar at a volume of 350 μL per well of the 24-well culture plate was found to be optimal for overnight incubation, without dehydrating the agar. Furthermore, the final purification step of phage isolation only yielded ~750 μL of phage suspended in SM buffer after passing through a 0.22 μm syringe filter. Thus, it was necessary to identify the minimal amount of phage suspension needed for host screening to avoid both consumption of the sample, and any additional phage propagation. Results showed that a volume using 2 μL of purified phages was sufficient to conduct miniaturized host range screening (Supplementary Table 1).

### 2.3. TAT of each assay

Comparisons of time required for each step within both assays were performed in triplicate using four *P. aeruginosa* isolates (three clinical isolates and one laboratory reference strain). Completion of the first step of host range screen consisting of the inoculation of overlay agar) of a single purified phage suspension against a single *P. aeruginosa* isolate required 77±2 minutes using the whole plate phage assay, while the 24-well phage assay required significantly less time with 16.5±2 minutes (Student’s t test; p<0.0001) (Figure 2a; Supplementary Table 2). Miniaturization of the host range screening saved approximately 60 minutes per phage sample against a single bacterial isolate. Extrapolation of the time saved when the breadth of host range was tested against 30 *P. aeruginosa* isolates resulted in a calculated reduction of 1,815 minutes (30.25 hours; 78.6%) when using the miniaturized assay.

The second step of the host range screen involves spot testing of phages on agar. It was found to require ~0.5 minutes (30 seconds; ±5 seconds) per phage sample per bacterial isolate using the whole plate phage assay, while the miniaturized 24-well phage assay required significantly less time at ~0.12 minutes (5 seconds; ±1 second) (Student’s t test; p<0.0016) (Figure 2b: Supplementary Table 2). Since host range screening was performed on 30 P. aeruginosa isolates in this study, ~11.0 minutes was calculated to be saved using the miniaturized assay.

**Figure 2.**
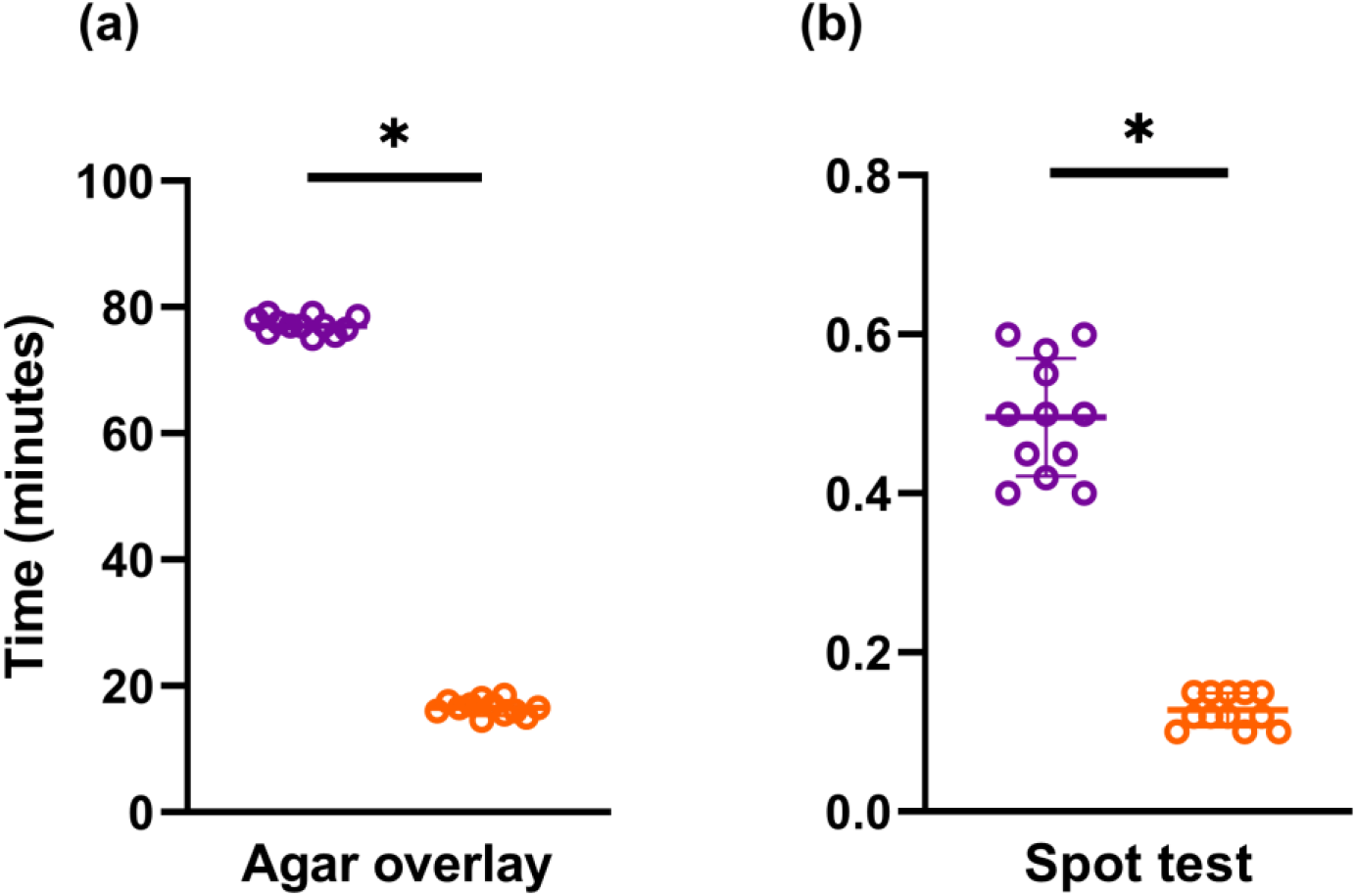
Comparison of the time required to screen a single phage suspension via whole plate phage assay (purple) or 24-well phage assay (orange). (a) The approximate amount of time needed to perform the inoculation of *P. aeruginosa* into molten overlay agar and pouring it onto either a LB agar plate for the whole plate phage assay or into an empty well for the 24-well phage assay. (b) Using pre-labelled and prepared phage suspension in the order of choice, the approximate time required to perform a spot test was observed to be less for the 24-well phage assay when compared to the whole plate phage assay. All data points as well as mean ± SD are shown from triplicate values obtained from the 4 isolates assessed.

### 2.4. Assay sensitivity and specificity

To assess assay specificity and sensitivity, phage suspensions (2 μL) were spotted onto both whole plate and 24-well phage assay plates in triplicate and visualized after 18 hours incubation at 37°C under aerobic conditions over different days. Data generated revealed that the miniaturized phage assay had a sensitivity and specificity score of 94.6% and 94.7% respectively (Table 1). The positive predictive value (PPV) was calculated at 86.4% and the negative predictive value (NPV) at 98% which indicated that for every 100 P. aeruginosa isolates tested, ~13 would potentially be a false positive while 2 might be a false negative.

**Table 1.**
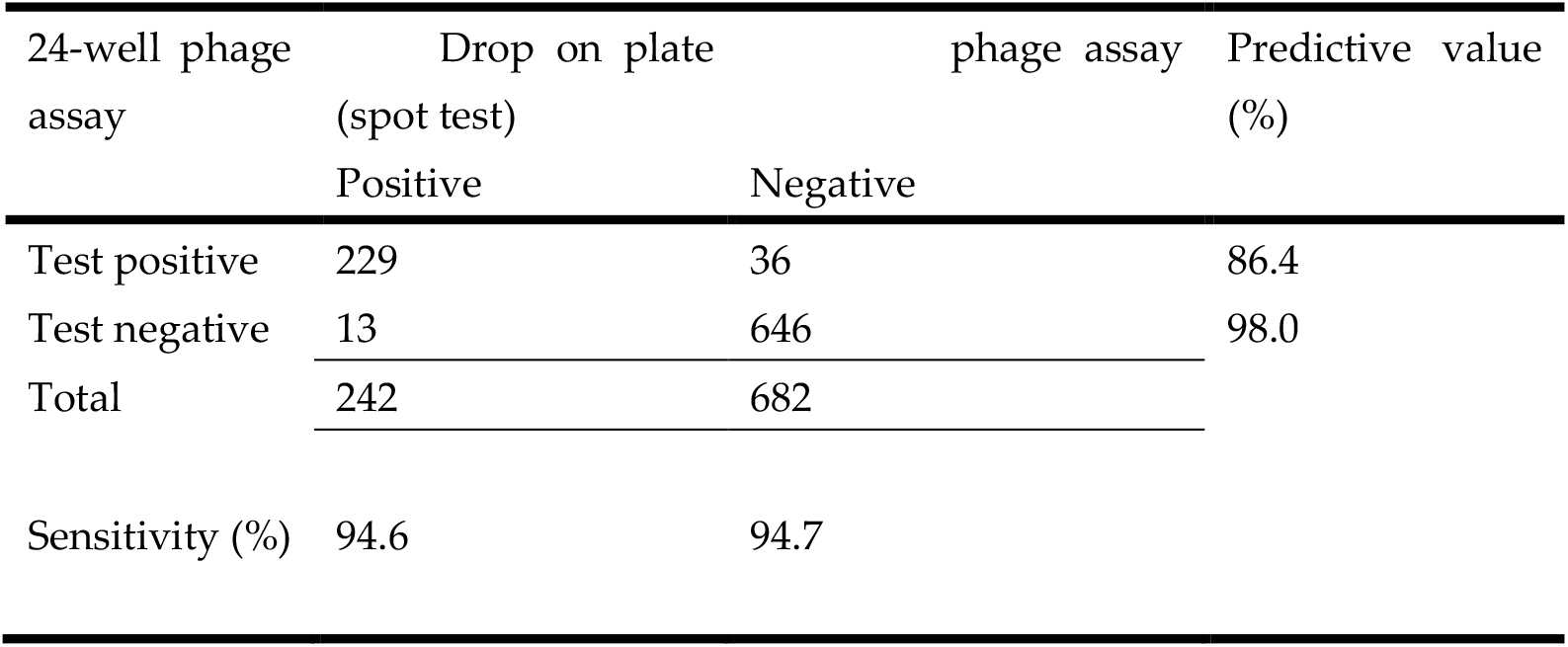
Sensitivity and specificity of 24-well phage assay

### 2.5. Assay sensitivity to low titers of phages

Sensitivities of the miniaturized 24-well phage assay to low titers of phages were then measured using a series of serially diluted phages. Both specificity and negative predictive value could not be calculated as the phages used in this experimental setup were measured against their respective propagating hosts, and hence no negative results were generated. The sensitivity of the miniaturized assay at low titers of phages was found to be 90.2%, which indicated that for every 100 *P. aeruginosa* isolates tested, ~9 may be a false positive.

### 2.6. Zones of clearance observable from whole plate vs. miniaturized 24-well phage assay

Plaque morphology was initially graded and classified into three categories: clear, opaque and no lysis from the whole plate phage assay. Further inspection of plaques revealed that both clear and opaque zones were present with varying morphologies. Firstly, completely clear plaques were observed and found to have zones of clearances that did not appear opaque (Figure 3a-b). Furthermore, within clear plaques, pinpoint colonies of bacteria could be observed following incubation (Figure 3a: yellow arrow). Secondly, opaque plaques were also seen and identified as having partial zones of clearance (Figure 3c) or a bullseye morphology (Figure 3d) with an accompanying ring-like formation.

**Figure 3.**
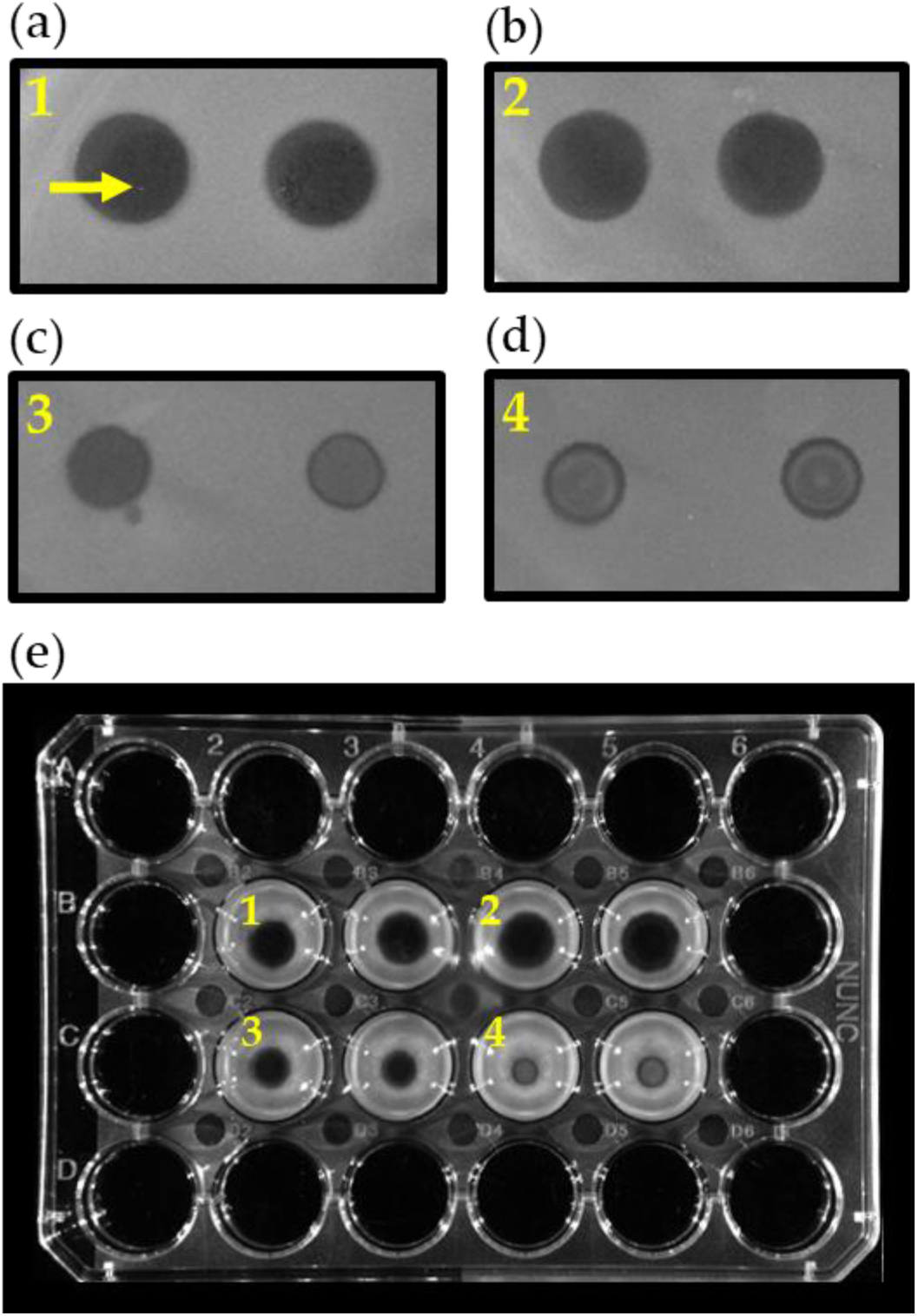
Visual inspection of zones of clearance demonstrated that morphology of the lytic activity were comparable between the whole plate phage assay and 24-well plate assay. The zones of clearance were not affected by the location of wells assigned nor modified based on the epi-light reflection and refraction when images were captured using the ChemiDoc XRS+ system. (a, b) Clear zones of lytic activity were observed on both the whole plate phage assay and 24-well phage assay, denoted by “1” and “2” respectively. (c) Turbid zones of clearance were observed on both methods, denoted by “3”. (e) Bullseye morphology of the plaques were observed on both the whole plate phage assay as well as the 24-well phage assay as denoted by “4”.

Using the developed miniaturized 24-well phage assay, plaque morphologies could also be visualized and classified with the same categories as the whole plate phage assay. The various plaques morphologies, namely clear and opaque, were also identified and comparable to that seen using the larger scale whole plate assay. Plaque morphologies of each plaque were also found not to be affected by the position of spot-test (Figure 3e) and overlay agar.

## 3. Discussion

Here, we describe a novel methodology to perform large scale testing of phage host range using inoculated overlay agar in a microtiter culture plate, which was found in our hands to be sensitive and accurate. Using 231 isolated phages propagated with 30 isolates of *P. aeruginosa*, we screened for their host range using both whole plate and 24-well miniaturized phage assays against four selected *P. aeruginosa* isolates. The miniaturized assay reported was found to be highly sensitive and specific at 94.6% and 94.7% respectively when compared to results derived from the whole plate phage assay. At the same time, the PPV of the 24-well phage assay when screened using all 231 phages against 30 *P. aeruginosa* isolates was 86.4% (Table 1) while 23 phages at low concentrations were 100% (Table 2). While this could be an inflation of PPV due to the higher prevalence of known positive populations, it highlights the usability of the screening tool described [35]. This is in line with the current process of host range screening using phages that were previously identified specific against the infective pathogens. Using various clinical isolates, this methodology was corroborated and could be used in the future to complement diagnostic tests. Furthermore, while this method utilized *P. aeruginosa* clinical isolates, it should be possible to adapt it, with modifications, to other bacteria including *Staphylococcus aureus*, which also causes respiratory infections in CF and has developed antibiotic resistance [36–39].

**Table 2.** Sensitivity and specificity of 24-well phage assay to low titers of phages

|  | 24-well phage assay | Drop on plate phage assay (spot test) |  | Predictive value (%) |
| --- | --- | --- | --- | --- |
|  |  | Positive (/23) | Negative (/23) |  |
| Test positive | 109 PFU/mL | 23 | 0 | 100 |
|  | 107 PFU/mL | 23 | 0 |  |
|  | 105 PFU/mL | 19 | 0 |  |
|  | 103 PFU/mL | 18 | 0 |  |
|  |  | 83 | 0 |  |
| Test negative | 109 PFU/mL | 0 | 0 |  |
|  | 107 PFU/mL | 0 | 0 |  |
|  | 105 PFU/mL | 4 | 0 |  |
|  | 103 PFU/mL | 5 | 0 |  |
|  |  | 9 | 0 |  |
| Total |  | 92 | 0 |  |
| Sensitivity (%) |  | 90.2 |  |  |

To complement diagnostic tests, the assay needs to be able to improve the TAT of current clinical laboratory standards. In a microbiology diagnostic laboratory, positive culture blood reporting timeline is routinely recorded. The results generated in approximately 72 hours consists of gram stains, pathogen identity and antimicrobial susceptibility test (AST) [40,41]. Specifically, in CF, sputum and swab samples are routinely collected during an active infection [42–44]. However, the TAT time for such cultures is not routinely recorded and may vary between laboratories due to differing local regulatory guidelines. Therefore, the miniaturized assay can be compared to the benchmark of a positive blood culture. While advancements were made to shorten the TAT from the initial flagging of positive blood cultures for both identification and AST [45–47], processes used to identify phage efficacy against the infective pathogen are still manual and time-consuming.

Phages can be isolated from a wide variety of sources, such as environmental or clinical samples, and may not be abundantly present. Therefore, the ability to screen for positive activity across a range of phage concentrations is both important and necessary. Throughout the purification process, phages are typically not propagated to a high titer. After the final purification step of phage isolation, titer still remains unknown and it is a time consuming process to perform subsequent titrations. While sensitive, our test was also specific, detecting as low as 10^3^ PFU/mL at 100% PPV (Table 2). Also, as shown by our results, we yielded only 750 μL of purified phages suspended in SM buffer after filtration. Thus it is important that our methodology was able to use both low concentrations of phages at a minimum volume to accommodate requirements to screen more bacterial isolates, which it was able to achieve easily.

While we have described a method that decreases the anticipated hours of manual labor significantly, further automation could also be applied. For example, larger volumes of phages could be utilized in the spot test step to cover the entire surface of the overlay agar, resulting in positive activity throughout the well. In addition, a spectrophotometer could be used to read the turbidity (optical density; OD) of the inoculated agar as an automated process. However, caution has to be exercised in the reliability of the results due to possible resistant colony growth in the middle of the clearing, giving rise to a false negative readout. Phage plaques and zones of clearing often display different morphologies, with some having a bullseye phenotype as identified in this study (Figure 3d). This could also result in a false negative result since spectrophotometers measures OD through the center of designated areas.

In order to enhance the morphology of phage plaques, tetrazolium salt dyes were used, however these are known to adversely affect phage kinetics and suppress plaque titers [48]. However, an automated microtiter plate reader has been described in literature using the OmniLog^™^ system to measure phage kinetics with the addition of tetrazolium salt dyes. The system utilizes the redox chemistry reaction to track the changes of bacterial viability after the application of phage formulations [49,50]. Using this chemical reaction, Adaptive Phage Therapeutics, Gaithersburg, MD have developed the Host Range Quick Test^™^ (HRQT^™^) under their sole proprietorship. While the HRQT^™^ had shown to be able to screen 5000 phages against one bacterial strain under 18 hours [51], this method may not be applicable to the majority of laboratories as it requires specific equipment that is not readily available. The HRQT^™^ also solely measures the kinetics of aerobic bacteria, but not anaerobes. However, our miniaturized phage assay could be applied to anaerobic phage screening as well. A limitation in our described methodology is the inability to track the emergence of resistant bacteria after the application of phages due to the test results being an endpoint readout. However, this could be overcome with a phage kill-curve experiment with varying titers after the identification of potential phages for therapeutic applications.

Overall, the assay has shown to be effective in reducing the TAT and manual labor hours in the laboratory. The miniaturized assay did not compromise both sensitivity and specificity at the reduction of time required. Although the sensitivity, specificity and predictive values could vary between research and diagnostic laboratories due to personnel and equipment differences, modifications could be performed to suit the workflow of laboratories. The usability and complementation of the 24-well phage assay screening methodology for therapeutic applications must be distinguished in the phage screening for therapeutic application process. In conclusion, we have demonstrated that miniaturization is the first step towards automation to test for efficacy and diversity of phages. Ultimately, based on high-throughput screening of phages against multiple pathogens, individual sequencing of bacteria and the application of machine learning could allow for in silico testing of individual pathogen susceptibility to catalogued libraries of phages.

## 4. Materials and Methods

### 4.1 Bacterial strains, culture conditions and inoculum preparations

All 29 clinical isolates of *P. aeruginosa* used were derived from individuals with cystic fibrosis unless otherwise stated. A laboratory reference strain of *P. aeruginosa* (PA01; ATCC 15692) was obtained from the University of Western Australia, Department of Microbiology Culture Collection (MCC) [52]. Overnight cultures of P. aeruginosa isolates were then propagated by inoculating a single colony of *P. aeruginosa* in LB Lennox broth (Becton Dickinson, USA) and incubating overnight at 37°C with orbital shaking (120 rpm).

### 4.2 Bacteriophage isolation and purification

Wastewater samples were collected from the Subiaco Wastewater Plant (Shenton Park, Western Australia, Australia), enriched, and screened for phages that were able to infect any member of the panel of 29 P. aeruginosa clinically-derived isolates (Supplementary Table 3) or PA01 [53,54]. Specifically, water samples were initially filtered through a 0.22 μm bottle-top filter (Nalgene^™^ Rapid-Flow^™^, ThermoFisher, USA), then supplemented with 1 M CaCl2 and 1 M MgCl2 to achieve a final concentration of 0.1 M CaCl2 and 0.1 M MgCl2. Double strength LB broth supplemented with a final concentration of 1 mM CaCl2 and 1 mM MgCl2 was added to the filtrate for 24-48 hours to enrich for phages. Enriched LB broth was then centrifuged at 4000 rpm for 10 minutes at room temperature (RT) and supernatants filtered through 0.22 μm syringe filters. Filtrates were then spot tested (drop-on-plate) on the *P. aeruginosa* isolates they were enriched with for lytic activity that had previously been line streaked on LB agar and air-dried for 15 minutes at RT. Five microliters of the filtrate was applied onto each streak and air-dried for another 15 minutes at RT. Agar plates were incubated under aerobic conditions at 37°C for 18 hours and phage presence determined via clearance visualization on the bacterial streaks. LB broth supplemented with 0.4% bacteriological agar (Becton Dickinson, USA) and 1 mM CaCl_2_ and 1 mM MgCl_2_ were routinely used as semi-solid media (overlay agar) [55]. Plaques were selected and purified in the propagating host via three purification rounds. Resulting stable phages (n= 231) were then propagated to a high titer and stored in SM buffer (100 mM NaCl, 8 mM Mg·SO_4_, 50 mM Tris-HCl, pH 7.5) at 4°C for further analysis.

### 4.3 Phage propagation to high titers

High titer phage stocks were propagated using solid media propagation as described [53]. Briefly, 100 μL of overnight culture was inoculated with 100 μL of purified phage suspension and incubated for 5 minutes at RT to allow adsorption of phages to the propagating host. Three milliliters of overlay agar being maintained at 55°C was added to the phage-host mixture and poured gently over LB agar plates. These were then incubated under aerobic conditions at 37°C for 24 hours, after which, plates were visually inspected for whole plate clearance. Five milliliters of SM buffer was then added to and evenly dispensed over the propagated plate and allowed to incubate for 15 minutes with gentle rocking at RT. Afterwards, the SM buffer was collected from the plate and filtered through 0.22 μm syringe filter. Phage titers were then enumerated using double agar overlay assay as described previously [55,56] and stored at 4°C.

### 4.4 Whole plate phage assay

Host range testing was performed using the gold standard whole plate phage assay, with modifications [55]. Here, 100 μL of overnight *P. aeruginosa* cultures were inoculated into 3 mL of molten overlay agar that was being maintained at 55°C. Inoculated overlay agars were then poured over LB agar plates, gently rocked to ensure an even spread and air-dried for 15 minutes at RT. Afterwards, 2 μL of purified phage suspension was dropped onto the agar and bacterial plates incubated for 18 hours at 37°C under aerobic conditions. Positive activity was then determined via visual confirmation of clearing and/ or plaque formation. Turbidity of zones of clearance were also assessed and graded (clear, opaque or no lysis). Negative activity was determined via no observable clearing or plaque formation. Phages with the broadest host range (active against ≥ 20/ 30 P. aeruginosa isolates) were serially diluted 10-fold from 10^9^ to 10^3^ PFU/mL and assayed (drop-on-plate) for plaque detection at 10^9^, 10^7^, 10^5^ and 10^3^ PFU/mL. Sterile SM buffer was spot tested onto overlay agar with and without P. aeruginosa as negative and media controls. Images of whole plate phage assay were captured using a ChemiDoc XRS+ imaging system using the epi-light filter (BioRad, USA).

### 4.5 Miniaturized phage assay

Assessment of host range against *P. aeruginosa* clinical isolates were repeated on a miniaturized scale, utilizing 24-well culture plates (Nunc^™^, ThermoFisher, USA). Briefly, 350 μL of overlay agar, inoculated with *P. aeruginosa*, were dispensed into each well of a 24-well culture plate and air-dried for 15 minutes at RT. Overnight cultures were inoculated proportionately to the amount needed to fill the required wells (100 μL of bacterial culture into 3 mL of molten overlay agar). Overlay agar with and without *P. aeruginosa* (350 μL) was also dispensed into an empty well on the 24-well culture plate to serve as negative and media controls. Two microliters of purified phage suspension was then added via droplet addition to assigned wells and plates were incubated overnight at 37°C under aerobic conditions. The following day, wells were inspected for positive or negative activity via manual visualization for zones of clearing or plaques and graded manually for the turbidity of zones of clearance (clear, opaque or no lysis). Serially diluted phages (10-fold increments: 109 to 103 PFU/mL) were then assayed (drop-on-plate) for plaque detection at 10^9^, 10^7^, 10^5^ and 10^3^ PFU/mL. Images of 24-well phage assay was captured using ChemiDoc XRS+ imaging system using the epi-light filter.

### 4.6 Statistics

Whole plate and miniaturized phage assays were performed in triplicate and replicated across 4 clinical isolates. All data points are presented as mean ± SD. Statistical analysis across methodologies of whole plate and miniaturized phage assays was performed with paired t-test, where a p-value < 0.05 was considered significant. (GraphPad Prism 9, USA).

## Supplementary Materials

The following are available online at www.mdpi.com/xxx/s1, Table S1: Optimization of media and phage volumes required for 24-well phage assay, Table S2: Breakdown of time required for individual steps of the host range screening for one bacterial isolate, Table S3: List of P. aeruginosa isolates used in the experimental setup.

## Author Contributions

Conceptualization, R.N.N. and A.K.; methodology, R.N.N. and A.K.; validation, R.N.N. and A.K.; formal analysis, R.N.N.; investigation, R.N.N.; resources, A.K.; data curation, R.N.N. and A.K.; writing—original draft preparation, R.N.N.; writing—review and editing, R.N.N., L.J.G., A.V., S.A.M., D.R.L., W.W.P.P., J.H., S.G.W., J.J.I., A.S.T., P.A.R., T.I., A.K., B.J.C. and S.M.S.; visualization, R.N.N.; supervision, A.K., B.J.C. and S.M.S.; project administration, R.N.N. and A.K.; funding acquisition, A.K. All authors have read and agreed to the published version of the manuscript.

## Institutional Review Board Statement

The study was conducted according to the guidelines of the Declaration of Helsinki and approved by the Royal Children’s Hospital Melbourne (approval no. HREC 25054). Written consent was obtained from the parents of the children enrolled into the study.

## Informed Consent Statement

Informed consent was obtained from all subjects involved in the study.

## Data Availability Statement

The data presented in this study are available within the article and supplementary document.

## Acknowledgments

We would like to sincerely thank Miss Kak Ming Ling for the initial input on the idea behind the manuscript. This article was written as part of the Australian Respiratory Early Surveillance Team for Cystic Fibrosis (AREST CF). The authors thank the subjects and families for their generous contributions to the program. Full membership of the AREST CF is available at www.arestcf.org.

## References

1. Serra-Burriel, M.; Keys, M.; Campillo-Artero, C.; Agodi, A.; Barchitta, M.; Gikas, A.; Palos, C.; López-Casasnovas, G. Impact of multi-drug resistant bacteria on economic and clinical outcomes of healthcare-associated infections in adults: Systematic review and meta-analysis. PLoS One 2020, 15, e0227139, doi:10.1371/journal.pone.0227139.

2. Huebner, C.; Roggelin, M.; Flessa, S. Economic burden of multidrug-resistant bacteria in nursing homes in Germany: a cost analysis based on empirical data. BMJ Open 2016, 6, e008458, doi:10.1136/bmjopen-2015-008458.

3. Knight, G.M.; McQuaid, C.F.; Dodd, P.J.; Houben, R.M.G.J. Global burden of latent multidrug-resistant tuberculosis: trends and estimates based on mathematical modelling. Lancet Infect. Dis. 2019, 19, 903–912, doi:10.1016/S1473-3099(19)30307-X.

4. Mave, V.; Chandanwale, A.; Kagal, A.; Khadse, S.; Kadam, D.; Bharadwaj, R.; Dohe, V.; Robinson, M.L.; Kinikar, A.; Joshi, S.; et al. High Burden of Antimicrobial Resistance and Mortality Among Adults and Children With Community-Onset Bacterial Infections in India. J. Infect. Dis. 2017, 215, 1312–1320, doi:10.1093/infdis/jix114.

5. Cystic Fibrosis Foundation 2018 PATIENT REGISTRY ANNUAL DATA REPORT; 2019;

6. Ruseckaite, R.; Ahern, S.; Ranger, T.; Dean, J.; Gardam, M.; Bell, S.; Burke, N. The Australian Cystic Fibrosis Data Registry Annual Report, 2017; 2019;

7. Smith, D.J.; Ramsay, K.A.; Yerkovich, S.T.; Reid, D.W.; Wainwright, C.E.; Grimwood, K.; Bell, S.C.; Kidd, T.J. Pseudomonas aeruginosa antibiotic resistance in Australian cystic fibrosis centres. Respirology 2016, 21, 329–337, doi:10.1111/resp.12714.

8. Schooley, R.T.; Biswas, B.; Gill, J.J.; Hernandez-Morales, A.; Lancaster, J.; Lessor, L.; Barr, J.J.; Reed, S.L.; Rohwer, F.; Benler, S.; et al. Development and use of personalized bacteriophage-based therapeutic cocktails to treat a patient with a disseminated resistant Acinetobacter baumannii infection. Antimicrob. Agents Chemother. 2017, 61, doi:10.1128/AAC.00954-17.

9. Aslam, S.; Courtwright, A.M.; Koval, C.; Lehman, S.M.; Morales, S.; Furr, C.L.L.; Rosas, F.; Brownstein, M.J.; Fackler, J.R.; Sisson, B.M.; et al. Early clinical experience of bacteriophage therapy in 3 lung transplant recipients. Am. J. Transplant. 2019, 19, 2631–2639, doi:10.1111/ajt.15503.

10. Dedrick, R.M.; Guerrero-Bustamante, C.A.; Garlena, R.A.; Russell, D.A.; Ford, K.; Harris, K.; Gilmour, K.C.; Soothill, J.; Jacobs-Sera, D.; Schooley, R.T.; et al. Engineered bacteriophages for treatment of a patient with a disseminated drug-resistant Mycobacterium abscessus. Nat. Med. 2019, 25, 730–733, doi:10.1038/s41591-019-0437-z.

11. Aslam, S.; Lampley, E.; Wooten, D.; Karris, M.; Benson, C.; Strathdee, S.; Schooley, R.T. Lessons Learned From the First 10 Consecutive Cases of Intravenous Bacteriophage Therapy to Treat Multidrug-Resistant Bacterial Infections at a Single Center in the United States. Open Forum Infect. Dis. 2020, 7, doi:10.1093/ofid/ofaa389.

12. Hoyle, N.; Zhvaniya, P.; Balarjishvili, N.; Bolkvadze, D.; Nadareishvili, L.; Nizharadze, D.; Wittmann, J.; Rohde, C.; Kutateladze, M. Phage therapy against Achromobacter xylosoxidans lung infection in a patient with cystic fibrosis: a case report. Res. Microbiol. 2018, 169, 540–542, doi:10.1016/j.resmic.2018.05.001.

13. LaVergne, S.; Hamilton, T.; Biswas, B.; Kumaraswamy, M.; Schooley, R.T.; Wooten, D. Phage Therapy for a Multidrug-Resistant Acinetobacter baumannii Craniectomy Site Infection. Open Forum Infect. Dis. 2018, 5, doi:10.1093/ofid/ofy064.

14. Zhvania, P.; Hoyle, N.S.; Nadareishvili, L.; Nizharadze, D.; Kutateladze, M. Phage therapy in a 16-year-old boy with netherton syndrome. Front. Med. 2017, 4, 1–5, doi:10.3389/fmed.2017.00094.

15. Chan, B.K.; Turner, P.E.; Kim, S.; Mojibian, H.R.; Elefteriades, J.A.; Narayan, D. Phage treatment of an aortic graft infected with Pseudomonas aeruginosa. Evol. Med. Public Heal. 2018, 2018, 60–66, doi:10.1093/emph/eoy005.

16. McCallin, S.; Alam Sarker, S.; Barretto, C.; Sultana, S.; Berger, B.; Huq, S.; Krause, L.; Bibiloni, R.; Schmitt, B.; Reuteler, G.; et al. Safety analysis of a Russian phage cocktail: From MetaGenomic analysis to oral application in healthy human subjects. Virology 2013, 443, 187–196, doi:10.1016/j.virol.2013.05.022.

17. Doub, J.B.; Ng, V.Y.; Johnson, A.J.; Slomka, M.; Fackler, J.; Horne, B.; Brownstein, M.J.; Henry, M.; Malagon, F.; Biswas, B. Salvage Bacteriophage Therapy for a Chronic MRSA Prosthetic Joint Infection. Antibiotics 2020, 9, 241, doi:10.3390/antibiotics9050241.

18. Law, N.; Logan, C.; Yung, G.; Furr, C.-L.L.; Lehman, S.M.; Morales, S.; Rosas, F.; Gaidamaka, A.; Bilinsky, I.; Grint, P.; et al. Successful adjunctive use of bacteriophage therapy for treatment of multidrug-resistant Pseudomonas aeruginosa infection in a cystic fibrosis patient. Infection 2019, 47, 665–668, doi:10.1007/s15010-019-01319-0.

19. Nir-Paz, R.; Gelman, D.; Khouri, A.; Sisson, B.M.; Fackler, J.; Alkalay-Oren, S.; Khalifa, L.; Rimon, A.; Yerushalmy, O.; Bader, R.; et al. Successful Treatment of Antibiotic-resistant, Poly-microbial Bone Infection With Bacteriophages and Antibiotics Combination. Clin. Infect. Dis. 2019, 69, 2015–2018, doi:10.1093/cid/ciz222.

20. Jikia, D.; Chkhaidze, N.; Imedashvili, E.; Mgaloblishvili, I.; Tsitlanadze, G.; Katsarava, R.; Morris, J.G.; Sulakvelidze, A. The use of a novel biodegradable preparation capable of the sustained release of bacteriophages and ciprofloxacin, in the complex treatment of multidrug-resistant Staphylococcus aureus-infected local radiation injuries caused by exposure to Sr90. Clin. Exp. Dermatol. 2005, 30, 23–26, doi:10.1111/j.1365-2230.2004.01600.x.

21. Chanishvili, N. Bacteriophages as Therapeutic and Prophylactic Means: Summary of the Soviet and Post Soviet Experiences. Curr. Drug Deliv. 2016, 13, 309–323, doi:10.2174/156720181303160520193946.

22. Kutateladze, M.; Adamia, R. Phage therapy experience at the Eliava Institute. Med. Mal. Infect. 2008, 38, 426–430, doi:10.1016/j.medmal.2008.06.023.

23. Kutateladze, M. Experience of the Eliava Institute in bacteriophage therapy. Virol. Sin. 2015, 30, 80–81, doi:10.1007/s12250-014-3557-0.

24. Chanishvili, N. Phage Therapy—History from Twort and d’Herelle Through Soviet Experience to Current Approaches. In Advances in Virus Research; 2012; pp. 3–40 ISBN 9780123944382.

25. Krestovnikova, V.A. Phage treatment and phage prophylactics and their approval in the works of the Soviet researchers. J. Microb. Epidemiol. Immun 1947, 3, 56–65.

26. Kokin, G.A. Use of bacteriophages in surgery. Sov. Med. (‘“Sovietskaya Meditsina”‘) 1941, 9, 15–18.

27. Nobrega, F.L.; Costa, A.R.; Kluskens, L.D.; Azeredo, J. Revisiting phage therapy: New applications for old resources. Trends Microbiol. 2015, 23, 185–191, doi:10.1016/j.tim.2015.01.006.

28. Merril, C.R.; Scholl, D.; Adhya, S.L. The prospect for bacteriophage therapy in Western medicine. Nat. Rev. Drug Discov. 2003, 2, 489–497, doi:10.1038/nrd1111.

29. Kutateladze, M.; Adamia, R. Bacteriophages as potential new therapeutics to replace or supplement antibiotics. Trends Biotechnol. 2010, 28, 591–595, doi:10.1016/j.tibtech.2010.08.001.

30. Holland, I.; Davies, J.A. Automation in the Life Science Research Laboratory. Front. Bioeng. Biotechnol. 2020, 8, doi:10.3389/fbioe.2020.571777.

31. Sarkozi, L.; Simson, E.; Ramanathan, L. The effects of total laboratory automation on the management of a clinical chemistry laboratory. Retrospective analysis of 36 years. Clin. Chim. Acta 2003, 329, 89–94, doi:10.1016/S0009-8981(03)00020-2.

32. Alshieban, S.; Al-Surimi, K. Reducing turnaround time of surgical pathology reports in pathology and laboratory medicine departments. BMJ Qual. Improv. Reports 2015, 4, u209223.w3773, doi:10.1136/bmjquality.u209223.w3773.

33. Dauwalder, O.; Landrieve, L.; Laurent, F.; de Montclos, M.; Vandenesch, F.; Lina, G. Does bacteriology laboratory automation reduce time to results and increase quality management? Clin. Microbiol. Infect. 2016, 22, 236–243, doi:10.1016/j.cmi.2015.10.037.

34. Miller, J.M.; Binnicker, M.J.; Campbell, S.; Carroll, K.C.; Chapin, K.C.; Gilligan, P.H.; Gonzalez, M.D.; Jerris, R.C.; Kehl, S.C.; Patel, R.; et al. A Guide to Utilization of the Microbiology Laboratory for Diagnosis of Infectious Diseases: 2018 Update by the Infectious Diseases Society of America and the American Society for Microbiologya. Clin. Infect. Dis. 2018, 67, e1–e94, doi:10.1093/cid/ciy381.

35. Vetter, T.R.; Schober, P.; Mascha, E.J. Diagnostic Testing and Decision-Making. Anesth. Analg. 2018, 127, 1085–1091, doi:10.1213/ANE.0000000000003698.

36. Schwerdt, M.; Neumann, C.; Schwartbeck, B.; Kampmeier, S.; Herzog, S.; Görlich, D.; Dübbers, A.; Große-Onnebrink, J.; Kessler, C.; Küster, P.; et al. Staphylococcus aureus in the airways of cystic fibrosis patients - A retrospective long-term study. Int. J. Med. Microbiol. 2018, 308, 631–639, doi:10.1016/j.ijmm.2018.02.003.

37. Wolter, D.J.; Emerson, J.C.; McNamara, S.; Buccat, A.M.; Qin, X.; Cochrane, E.; Houston, L.S.; Rogers, G.B.; Marsh, P.; Prehar, K.; et al. Staphylococcus aureus Small-Colony Variants Are Independently Associated With Worse Lung Disease in Children With Cystic Fibrosis. Clin. Infect. Dis. 2013, 57, 384–391, doi:10.1093/cid/cit270.

38. Steinig, E.J.; Duchene, S.; Robinson, D.A.; Monecke, S.; Yokoyama, M.; Laabei, M.; Slickers, P.; Andersson, P.; Williamson, D.; Kearns, A.; et al. Evolution and Global Transmission of a Multidrug-Resistant, Community-Associated Methicillin-Resistant Staphylococcus aureus Lineage from the Indian Subcontinent. MBio 2019, 10, doi:10.1128/mBio.01105-19.

39. Dasenbrook, E.C.; Checkley, W.; Merlo, C.A.; Konstan, M.W.; Lechtzin, N.; Boyle, M.P. Association between respiratory tract methicillin-resistant Staphylococcus aureus and survival in cystic fibrosis. JAMA-J. Am. Med. Assoc. 2010, 303, 2386–2393, doi:10.1001/jama.2010.791.

40. Tabak, Y.P.; Vankeepuram, L.; Ye, G.; Jeffers, K.; Gupta, V.; Murray, P.R. Blood Culture Turnaround Time in U.S. Acute Care Hospitals and Implications for Laboratory Process Optimization. J. Clin. Microbiol. 2018, 56, 500–518, doi:10.1128/JCM.00500-18.

41. Winter, A.; Jones, W.S.; Allen, A.J.; Price, D.A.; Rostron, A.; Filieri, R.; Graziadio, S. The Clinical Need for New Diagnostics in the Identification and Management of Patients with Suspected Sepsis in UK NHS Hospitals: A Survey of Healthcare Professionals. Antibiotics 2020, 9, 737, doi:10.3390/antibiotics9110737.

42. Ronchetti, K.; Tame, J.-D.; Paisey, C.; Thia, L.P.; Doull, I.; Howe, R.; Mahenthiralingam, E.; Forton, J.T. The CF-Sputum Induction Trial (CF-SpIT) to assess lower airway bacterial sampling in young children with cystic fibrosis: a prospective internally controlled interventional trial. Lancet Respir. Med. 2018, 6, 461–471, doi:10.1016/S2213-2600(18)30171-1.

43. Mussaffi, H.; Fireman, E.M.; Mei-Zahav, M.; Prais, D.; Blau, H. Induced sputum in the very young: A new key to infection and inflammation. Chest 2008, 133, 176–182, doi:10.1378/chest.07-2259.

44. Equi, A.C. Use of cough swabs in a cystic fibrosis clinic. Arch. Dis. Child. 2001, 85, 438–439, doi:10.1136/adc.85.5.438.

45. Charnot-Katsikas, A.; Tesic, V.; Love, N.; Hill, B.; Bethel, C.; Boonlayangoor, S.; Beavis, K.G. Use of the Accelerate Pheno System for Identification and Antimicrobial Susceptibility Testing of Pathogens in Positive Blood Cultures and Impact on Time to Results and Workflow. J. Clin. Microbiol. 2017, 56, doi:10.1128/JCM.01166-17.

46. Schoepp, N.G.; Schlappi, T.S.; Curtis, M.S.; Butkovich, S.S.; Miller, S.; Humphries, R.M.; Ismagilov, R.F. Rapid pathogen-specific phenotypic antibiotic susceptibility testing using digital LAMP quantification in clinical samples. Sci. Transl. Med. 2017, 9, eaal3693, doi:10.1126/scitranslmed.aal3693.

47. Idelevich, E.A.; Storck, L.M.; Sparbier, K.; Drews, O.; Kostrzewa, M.; Becker, K. Rapid Direct Susceptibility Testing from Positive Blood Cultures by the Matrix-Assisted Laser Desorption Ionization–Time of Flight Mass Spectrometry-Based Direct-on-Target Microdroplet Growth Assay. J. Clin. Microbiol. 2018, 56, doi:10.1128/JCM.00913-18.

48. Hurst, C.J.; Blannon, J.C.; Hardaway, R.L.; Jackson, W.C. Differential effect of tetrazolium dyes upon bacteriophage plaque assay titers. Appl. Environ. Microbiol. 1994, 60, 3462–3465, doi:10.1128/aem.60.9.3462-3465.1994.

49. Estrella, L.A.; Quinones, J.; Henry, M.; Hannah, R.M.; Pope, R.K.; Hamilton, T.; Teneza-mora, N.; Hall, E.; Biswajit, B. Characterization of novel Staphylococcus aureus lytic phage and defining their combinatorial virulence using the OmniLog^®^ system. Bacteriophage 2016, 6, e1219440, doi:10.1080/21597081.2016.1219440.

50. Henry, M.; Biswas, B.; Vincent, L.; Mokashi, V.; Schuch, R.; Bishop-Lilly, K.A.; Sozhamannan, S. Development of a high throughput assay for indirectly measuring phage growth using the OmniLog TM system. Bacteriophage 2012, 2, 159–167, doi:10.4161/bact.21440.

51. Caflisch, K.M.; Patel, R. Implications of Bacteriophage-and Bacteriophage Component-Based Therapies for the Clinical Microbiology Laboratory. J. Clin. Microbiol. 2019, 57, doi:10.1128/JCM.00229-19.

52. Dunn, N.W.; Holloway, B.W. Pleiotropy of p-fluorophenylalanine-resistant and antibiotic hypersensitive mutants of Pseudomonas aeruginosa. Genet. Res. 1971, 18, 185–197, doi:10.1017/S0016672300012593.

53. Bonilla, N.; Rojas, M.I.; Netto Flores Cruz, G.; Hung, S.; Rohwer, F.; Barr, J.J. Phage on tap–a quick and efficient protocol for the preparation of bacteriophage laboratory stocks. PeerJ 2016, 4, e2261, doi:10.7717/peerj.2261.

54. Van Twest, R.; Kropinski, A.M. Bacteriophage enrichment from water and soil. Methods Mol. Biol. 2009, 501, 15–21, doi:10.1007/978-1-60327-164-6_2.

55. Kropinski, A.M.; Mazzocco, A.; Waddell, T.E.; Lingohr, E.; Johnson, R.P. Enumeration of Bacteriophages by Double Agar Overlay Plaque Assay. In Bacteriophages: Methods and Protocols; Clokie, M.R.J., Kropinski, A.M., Eds.; Methods in Molecular Biology; Humana Press: Totowa, NJ, 2009; Vol. 501, pp. 69–76 ISBN 978-1-58829-682-5.

56. Abedon, S.T.; Katsaounis, T.I. Basic phage mathematics. In Methods in Molecular Biology; 2018; Vol. 1681, pp. 3–30 ISBN 9781493973439.

