## Supplementary Table 1 for "Development and validation of a miniaturized host range screening assay for bacteriophages"

**Supplementary Table 1.** Optimization of media and phage volumes required for 24-well phage assay.

| **Overlay agar (µL)** | **Phage suspension (µL)** | **Length of incubation (hours)** | **Results (+/-)** |
| --- | --- | --- | --- |
| 200 | 2 | 18 | - |
|  | 3 | 18 | - |
|  | 4 | 18 | - |
|  | 5 | 18 | - |
| 250 | 2 | 18 | - |
|  | 3 | 18 | - |
|  | 4 | 18 | - |
|  | 5 | 18 | - |
| 300 | 2 | 18 | + |
|  | 3 | 18 | + |
|  | 4 | 18 | + |
|  | 5 | 18 | + |
| 350 | 2 | 18 | + |
|  | 3 | 18 | + |
|  | 4 | 18 | + |
|  | 5 | 18 | + |
| 400 | 2 | 18 | + |
|  | 3 | 18 | + |
|  | 4 | 18 | + |
|  | 5 | 18 | + |
| 450 | 2 | 18 | + |
|  | 3 | 18 | + |
|  | 4 | 18 | + |
|  | 5 | 18 | + |
| 500 | 2 | 18 | + |
|  | 3 | 18 | + |
|  | 4 | 18 | + |
|  | 5 | 18 | + |
