## Supplementary Table 2 for "Development and validation of a miniaturized host range screening assay for bacteriophages"

**Supplementary Table 2.** Breakdown of time required for individual steps of the host range screening for one bacterial isolate.

|  |  | **Time required per step (minutes)** | **Total time required (minutes)** |
| --- | --- | --- | --- |
| **Whole plate phage assay** |  |  |  |
| LB agar plate | Pour agar plates | 1 |  |
|  | Allow the plates to dry | 60 |  |
| Overlay agar | Inoculation of *P. aeruginosa* | 0.5 |  |
|  | Overlay inoculated molten agar | 0.5 |  |
| Dry the plates in the hood |  | 15 |  |
|  |  |  | 77 |
| Spot test of phages |  | 0.5 |  |
|  |  |  | 0.5 |
| **24-well phage assay** |  |  |  |
| Overlay agar | Inoculation of *P. aeruginosa* | 1 |  |
|  | Dispense inoculated molten agar | 0.5 |  |
| Dry the 24-well plates in the hood |  | 15 |  |
|  |  |  | 16.5 |
| Spot test of phages |  | 0.12* |  |
|  |  |  | 0.6 |

* The amount of time required to perform a spot test for one bacterial isolate was derived from the equal distribution of time used to spot 24 phage and/or buffer suspensions onto each corresponding well using a multichannel pipette.
