## Supplementary Table 3 for "Development and validation of a miniaturized host range screening assay for bacteriophages"

**Supplementary Table 3.** List of ARESTCF *P. aeruginosa* isolates used in the experimental setup.

| **M1C** | **Sample source** | **Colony morphology** |
| --- | --- | --- |
| 79 | Sputum | Rough |
| 64 | BAL | Rough |
| 54 | Sputum | Rough |
| 50 | BAL | Rough |
| 123 | Sputum | Rough |
| 77 | BAL | Rough |
| 10 | Sputum | Rough |
| 34 | Sputum | Rough |
| 14 | Sputum | Rough |
| 66 | BAL | Rough |
| 100 | Sputum | Rough |
| 131 | Sputum | Rough |
| 141 | CS | Rough |
| 39 | Sputum | Rough |
| 67 | Sputum | Rough |
| 90 | BAL | Rough |
| 124 | BAL | Rough |
| 130 | Sputum | Rough |
| 94 | BAL | Rough |
| 108 | Sputum | Rough |
| 81 | BAL | Rough |
| 74 | CS | Rough |
| 58 | CS | Rough |
| 37 | CS | Rough |
| 73 | BAL | Rough |
| 52 | BAL | Rough |
| 86 | BAL | Rough |
| 4 | Sputum | Mucoid |
| 93 | Sputum | Mucoid |

Abbreviations: BAL – Bronchoalveolar Lavage; CS – Cough Swabs
